# Testing a general theory for flowering time shift as a function of growing season length

**DOI:** 10.1101/2025.02.05.636594

**Authors:** John S. Park, John Jackson, Anna Bergsten, Jon Ågren

**Affiliations:** Department of Biology, University of Oxford, Oxford, United Kingdom; Department of Conservation Biology and Global Change, Estación Biológica de Doñana (EBD-CSIC), Sevilla, Spain; Plant Ecology and Evolution, Department of Ecology and Genetics, Evolutionary Biology Centre, Uppsala University, Uppsala, Sweden

**Keywords:** phenology, life history theory, flowering time, growing season length, perennial plants

## Abstract

Understanding the effects of climate change on the evolution of phenological timing, such as start of flowering, is of major interest because phenology is critical for fitness of populations, and underpins many ecological dynamics. Much recent research has focused on the correlation between phenological timing and the arrival of spring. However, the evolutionarily optimal seasonal timing should depend also on the duration of the growing season, within which entire annual life cycles must unfold. Optimal energy allocation theory can explicitly address life-history scheduling in a seasonal environment, and be used to predict how scheduling should adaptively respond to seasonality shifts. Here we extend a seminal theoretical framework for perennial plant scheduling by Iwasa & Cohen (1989), and predict a specific nonlinear relationship between growing season length and optimal flowering time expressed as number of days after the start of the growing season. We tested and found strong support for this *a priori* prediction in two independent common-garden experiments with purple loosestrife (*Lythrum salicaria*) and European goldenrod (*Solidago virgaurea*) populations sampled along latitudinal gradients in Sweden. Climate warming is commonly associated with changes in both the start and the duration of the growing season. Considering both effects, our findings suggest that as springs start earlier and growing seasons lengthen, shifts in optimal flowering time expressed as calendar date may initially stall before accelerating, potentially explaining observed variation in phenological shifts across systems. More broadly, we show how mechanistic life history theory can advance understanding of phenological change beyond correlative conclusions.

**SIGNIFICANCE STATEMENT:** The optimal timing for a plant to flower depends both on when spring begins and on the total length of the growing season—both shifting with climate change. However, the adaptation of flowering time to changes in growing season length is much less explored than to an advancing spring. Building on existing theory, we predict a nonlinear relationship between growing season length and optimal flowering time. Data on two species (purple loosestrife and European goldenrod) grown in common gardens strongly support this prediction. Our results demonstrate why species can vary in their observed phenological responses to climate change. More broadly, we highlight the power of mechanistic life history theory for explaining and predicting phenological shifts.

## INTRODUCTION

When to reproduce is a critical fitness-determining decision for seasonal organisms, confined to a limited window of favorable conditions: the growing season. Climate change is both advancing the onset and prolonging the duration of this window in many systems (1, 2). These perturbations have been associated with global changes in biological timing known as phenological shifts (3, 4). While in most cases phenology advances, the magnitude and direction of shifts vary considerably among individual organisms, populations, and species (5–10). To understand the impacts of climate change, it is therefore important to explain the causes and shapes of variation in phenological responses to changes in the onset and duration of seasons.

Observed phenology, such as flowering time, results from the complex coordination of many interdependent life-history processes that include resource acquisition and allocation to growth, reproduction and survival. These processes typically trade off or covary, each with its own time requirements and contribution to fitness (11–14). Life history theory explains how natural selection for optimal combinations of such processes, within biological constraints, produces the great diversity of strategies seen in nature (15). While phenology is widely recognized as an outcome of life-history processes, modern analyses often overlook how interdependent life-history processes combine to shape observed phenological patterns and their changes (7, 11, 16, 17).

Changes in growing season length can alter the consequences of life history trade-offs, requiring adjustments to annual life-history schedules to maximize long-term fitness. Predicting evolutionarily optimal adjustments can be mathematically complex because life-history processes are bound by invariable temporal order (18) and trade-offs can be nonlinear and multidimensional. For perennial organisms, an added layer of complexity is balancing annual scheduling with optimal resource storage for subsequent years to maximize long-term fitness. Therefore, life history trade-offs traverse intra- and interannual timescales (17). Yet, most empirical investigations have focused on how system-specific cues, such as spring onset, proximately induce isolated life cycle events (7, 17). How changes in the total length of the growing season—the template within which entire annual life cycles must unfold—shape phenological schedules is far less explored. While the relevant seasonal limits (*e.g.,* temperature or precipitation thresholds) vary by system, limited time is a universal constraint in seasonal systems, offering a unifying perspective on phenological adaptation to seasonality.

Optimal energy allocation models provide a direct mathematical translation of the annual life-history scheduling problem in a general and species agnostic manner (19, 20). Yet, they have been surprisingly overlooked in the recent surge of empirical work. These models address how an individual should continuously assimilate and expend energy across various life-history activities to maximize fitness, considering the environmental context and dynamic consequences of decisions through time. Optimal strategies have been used to analyze population- and species-level differentiation, from the perspective of long-term selection (19, 21). Classic optimal energy allocation models were designed to explore broader life-history divergence across taxa, *e.g.*, itero/semelparity (22), age at first reproduction (22, 23), and in/determinate growth (24). These classic models typically use the year (age) as the basic unit of time. However, using continuous-time functions that can reflect and be optimized for arbitrarily fine time-scale processes, a small group of theorists in the 1970’s and 80’s explored adaptive optimality of *within-year* life-history schedules (20, 21, 25–28). For multi-year life histories, such models use dynamic programming (19) to incorporate how a given year’s optimal solution is adjusted by its consequences for years later in life.

In a seminal paper (29), Iwasa & Cohen provided a framework for the optimal annual scheduling of vegetative and reproductive activity of perennial plants. Their work builds on that of Schaffer (30) and others (20, 22, 26, 31). To introduce what follows, we summarize the original Iwasa-Cohen model in Fig. 1 and Box 1 (for a more general review of optimal energy allocation models, see (19, 21)). Here we trace a particular derivation in the Iwasa-Cohen model regarding how growing season length controls optimal flowering time.

**Figure 1.**
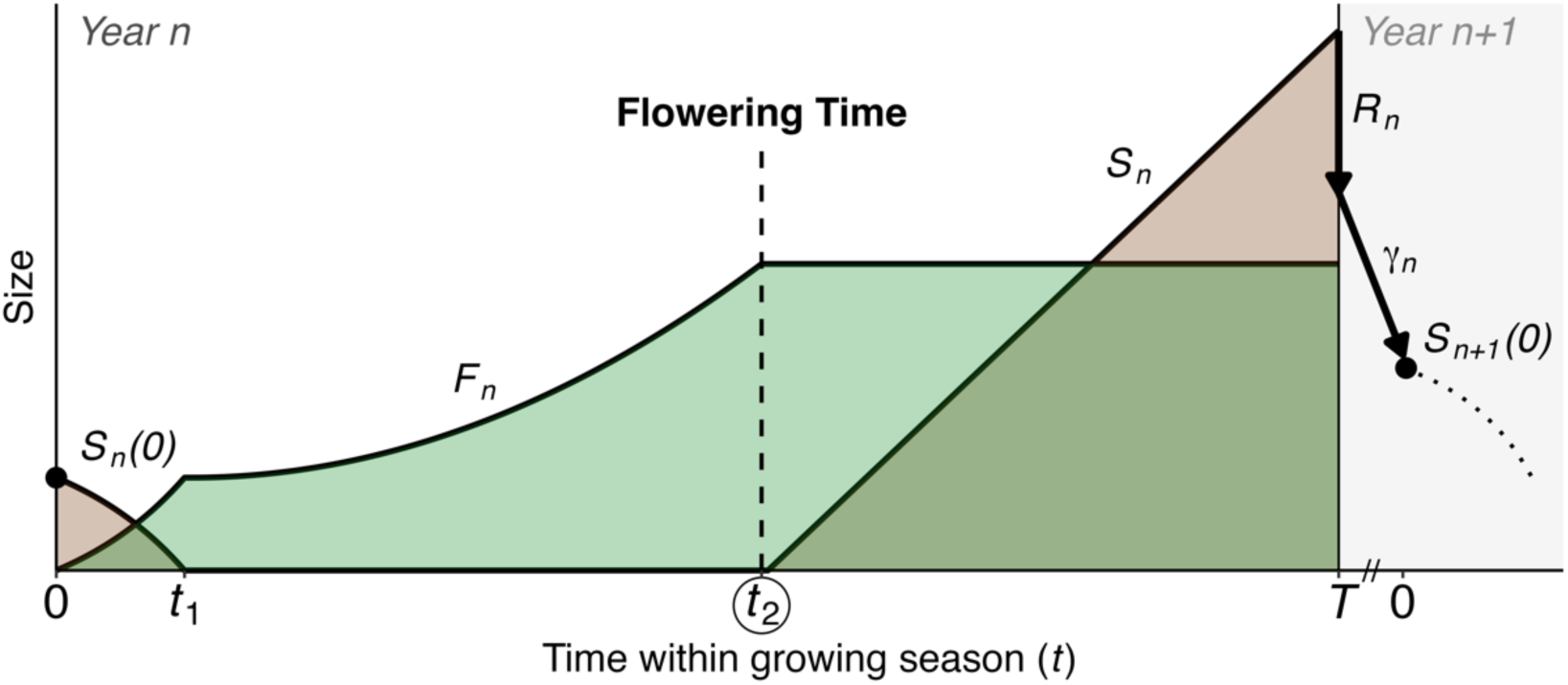
Schematic illustration of the annual schedule of a perennial plant adapted from Iwasa & Cohen (1989). The origin 𝑡 = 0 is the beginning of the growing season, not the year, and thus time points along 𝑡 indicate the number of days relative to the start of the growing season. Two curves with colored areas show growth functions of the “production” (𝐹_𝑛_) and “storage” (𝑆_𝑛_) parts in year 𝑛. “Storage” in the original model includes both actual storage for future use as well as reproductive activity. 𝐹_𝑛_ is discarded at the end of growing season 𝑇, and 𝑆_𝑛_(𝑇) is partially allocated to year 𝑛’s reproductive investment 𝑅_𝑛_, and partially lost due to imperfect storage efficiency 𝛾. The storage size to start the following year’s life cycle is thus 𝑆_𝑛+1_(0) = 𝛾[𝑆_𝑛_(𝑇) − 𝑅_𝑛_]. Discontinuity marks on the x-axis indicate time between growing seasons of year 𝑛 and year 𝑛 + 1 (*i.e.,* winter). Small 𝑡’s on the x-axis indicate the two critical switching points, with 𝑡_2_ being the beginning of reproductive investment, *i.e.* flowering time, the focus of our study.

Specifically, we examine Iwasa & Cohen’s solution for optimal flowering time for polycarpic perennial plants that start growth each year with resources retained from the preceding year:

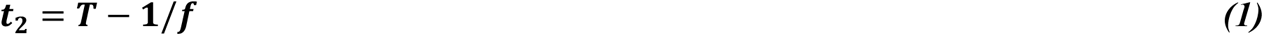

where 𝑡_2_ is optimal flowering time counting from the start of the growing season (not the start of the year), 𝑇 is the length of the growing season, and 𝑓 is the maximum relative growth rate (g/g/day), *i.e.,* relative growth when the plant is small. It is important to note that this deceptively simple prediction is an outcome of optimizing 28 initial parameters that mathematically cancelled out, rather than being disregarded (29). Despite its simplicity and broad implications for phenology, this prediction has gone largely unexplored and has not been empirically tested to our knowledge.

Note that relative growth rate 𝑓, the mediating parameter, is a key plant life-history trait (15, 32–34) that varies clinally among populations with latitude, thermal regimes, and growing season length (35–40). Although local factors such as herbivory can impact optimal growth rate (*e.g.,* (41)), general life history theory predicts that shorter growing seasons should select for increased relative growth rates so that development can be completed within the available time. This prediction has been repeatedly supported empirically (42). Hence, to test the Iwasa-Cohen prediction for perennial plant life histories across a range of growing season lengths, we adjust Eq. 1 to:

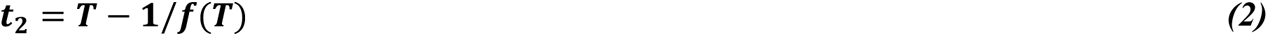

where 𝑓(𝑇) = 𝑎⁄1 + 𝑒^−𝑏(𝑇+𝑐)^is a logistic function of relative growth rate dependent on growing season length, with three parameters: 𝑎 is the upper bound of 𝑓, 𝑏 is the rate of change (steepness), and 𝑐 is a horizontal transformation parameter. This extension generates a curve with a long initial phase of steady increase, followed by a slowdown, then a fall in optimal flowering time measured from the start of the growing season (Fig. 2). The nonlinear shape of 𝑡_2_(𝑇) has two important biological implications. First, it suggests a potential shift in the rate of change in optimal flowering time, measured from the start of the season, as growing season expands. Second, if two populations or species occur on different realms of 𝑇, or have different functions of 𝑓(𝑇), their changes in optimal flowering time can be quite different given the same absolute change in 𝑇.

**Figure 2.**
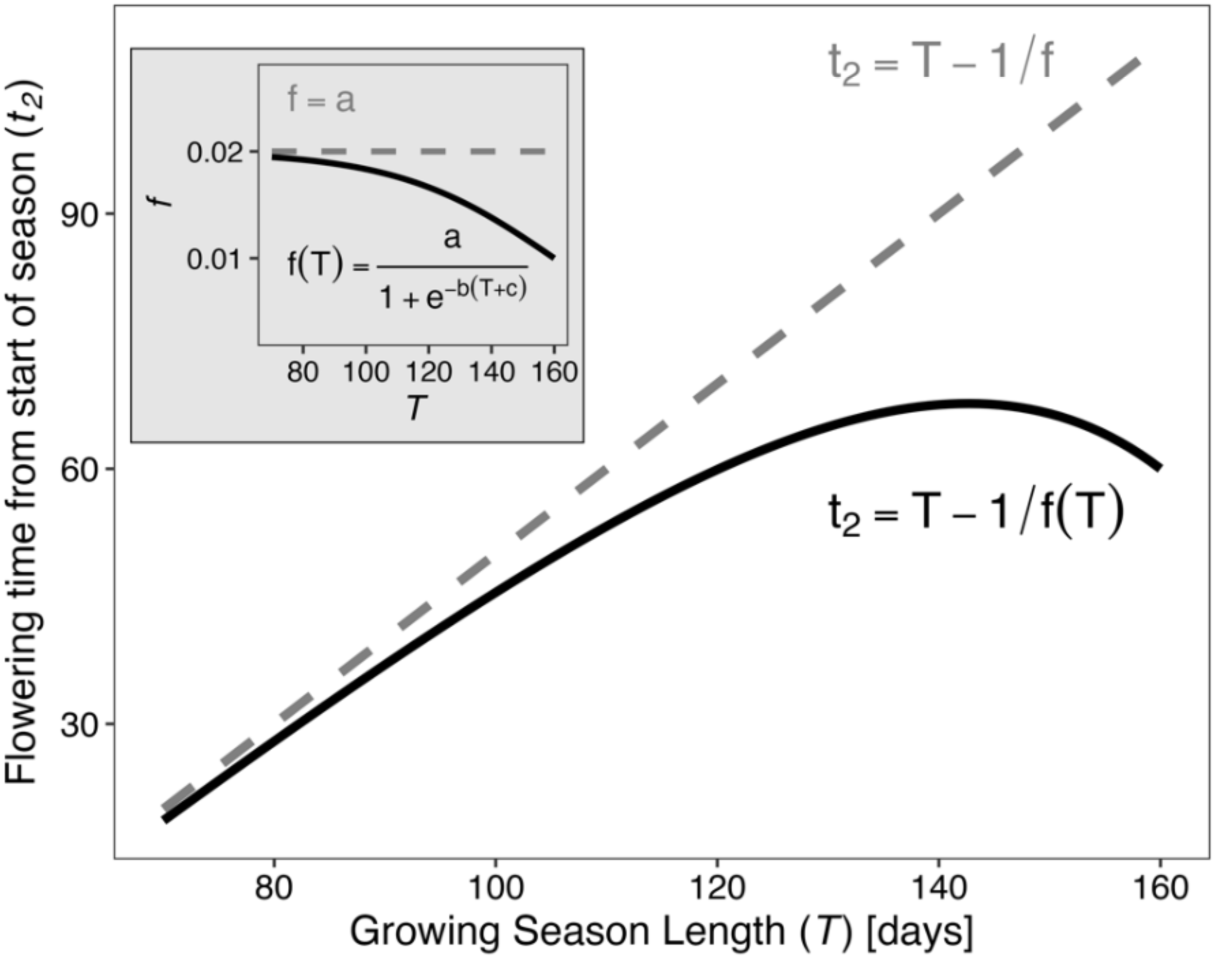
Modification of the Iwasa-Cohen solution (dashed lines) to incorporate clinal variation in maximum relative growth rate (solid lines), illustrating the nonlinear change in flowering time, expressed as time since beginning of the growing season. Inset shows functions for *f* underlying flowering time (𝑡_2_) in main plot. Parameter values for illustration: a = 0.02, b = -0.04, c = -160.

To test this prediction, we take advantage of the natural gradient of growing season length across latitudes (SI Appendix Fig. S1). We used data from two independent common-garden experiments with plants sourced from populations across broad latitudinal (57.4° to 68.4°) gradients in Sweden. The first comprised 15 populations of the perennial herb *Lythrum salicaria* (43). The second comprised 8 boreal populations of the perennial herb *Solidago virgaurea*. We used historical (1961-) daily weather data to calculate the mean 𝑇 at each source site. Then, we tested whether flowering time (measured as time since beginning of the growing season) varies following the predicted curvature (Fig. 2), given the growing season length in each source population. Finally, since source populations have varying start dates of the season *in situ*, we combined the estimated impact of length and the start of the growing season to estimate *in situ* optimal flowering time expressed as day of the year. Using a space-for-time substitution, the concurrent effect of earlier springs and longer seasons also reflects a prediction of optimal flowering date change under climate change at a given site.

## RESULTS

### Flowering time in both species follows the predicted nonlinear link to growing season length

Mean growing season lengths in *Lythrum* source population sites ranged from 134.8 to 205.4 days, and in *Solidago* populations from 116.7 to 159.5 days. In the common-garden experiments, population means of flowering time spanned 21 days for *Lythrum* (18 Aug-8 September) and 21 days for *Solidago* (23 Aug-13 September) (full individual data shown in SI Appendix Fig. S1a). Posterior estimates for fitted shape parameters in 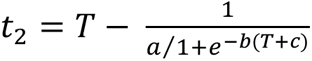 were 𝑎 = 0.021 [99% credible interval: 0.018, 0.024] , b = −0.010 [−0.012, −0.009] , and c = −144 [−168, −120] across *Lythrum* populations, and a = 0.026 [0.022, 0.039] , b = −0.043 [−0.083, −0.0.023], and c = −162[−167, −138] across *Solidago* populations (Fig. 3) (SI Appendix Text S1 for full model structure). In both species, we found strong statistical support for our modified Iwasa-Cohen model that predicted a specific nonlinear functional relationship between growing season length and optimal flowering time. Using leave-one-out cross validation, we found that the modified model substantially increased predictive performance compared to models that assumed no variation in the mediating parameter 𝑓 across 𝑇 (Table 1).

**Figure 3.**
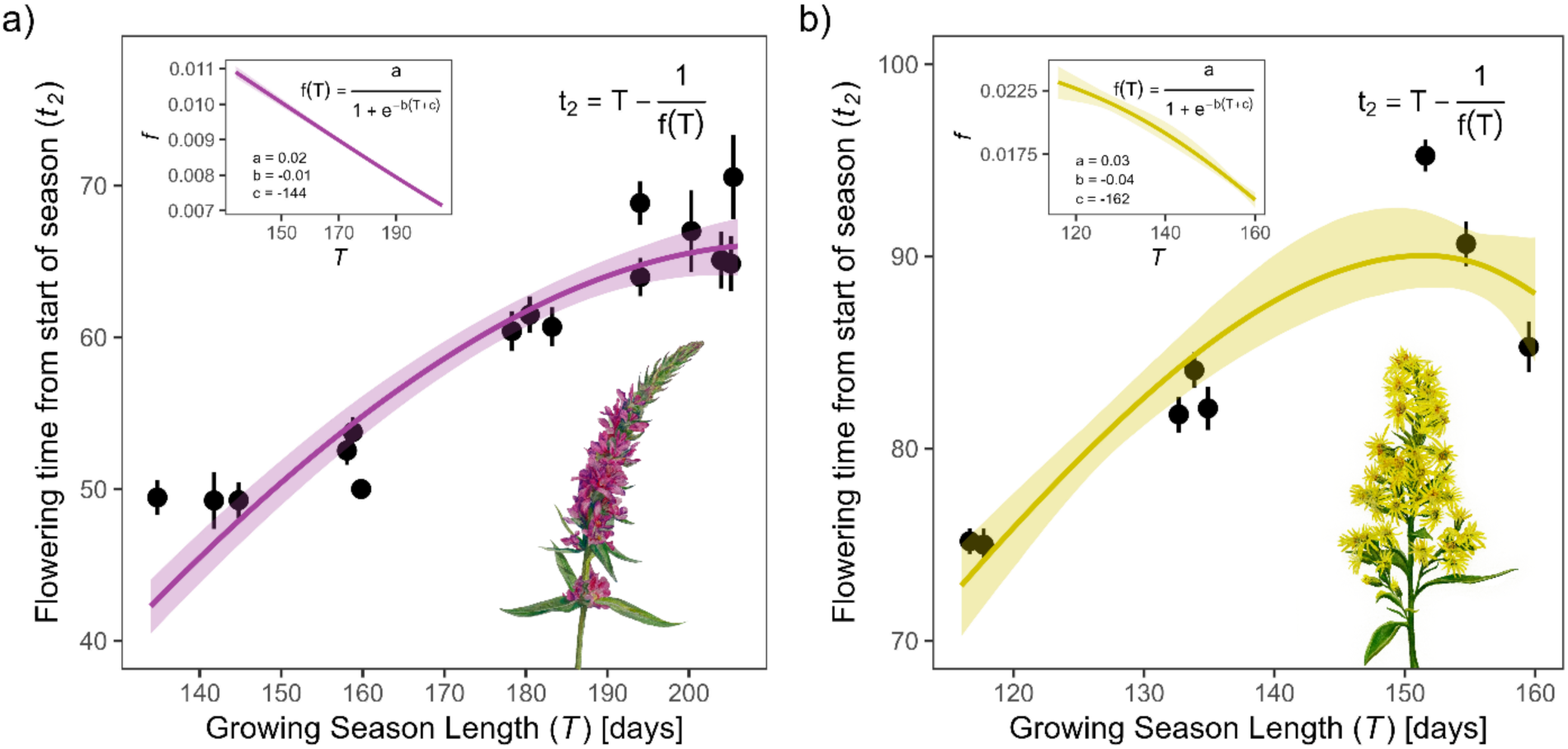
The fitted model of flowering time as a function of growing season length (main plots), and corresponding function of maximum relative growth rate 𝑓(𝑇) (insets), for a) *Lythrum* and b) *Solidago* common-garden experiment data. Points are population means of flowering time, and bars are ± SE. Shaded ribbons in both insets and main plots indicate posterior credible/prediction intervals at the 99% level. Prediction intervals for 𝑡_2_ include parameter uncertainty and posterior mean population level variance in 𝑡_2_, 𝜎 (SI Appendix Text S1). *Illustrations courtesy of Becca Huggins*.

**Table 1.** Predictive performance and leave-one-out information criterion (LOOIC) comparison between the modified, nonlinear Iwasa-Cohen model (𝑡_2_ = 𝑇 − 1/𝑓(𝑇)) and a model with constant 𝑓 (𝑡_2_ = 𝑇 − 1/𝑓). *elpd* is the expected log pointwise predictive density, Δ *elpd* gives the difference in *elpd* between models, and *elpd se* indicates the standard error in *elpd*.

| Lythrum model | elpd | Δelpd | elpd se | LOOIC |
| --- | --- | --- | --- | --- |
| Iwasa-Cohen model [f(T)] | -1,821.2 | 0.0 | 30.9 | 3,642.5 |
| Constant f | -2,172.3 | -351.1 | 15.8 | 4,344.6 |
| Solidago model | elpd | Δelpd | elpd se | LOOIC |
| Iwasa-Cohen model [f(T)] | -583.8 | 0.0 | 12.6 | 1,167.5 |
| Constant f | -716.1 | -132.3 | 8.9 | 1,432.2 |

The increasing curvature of 𝑡_2_ = 𝑇 − 1/𝑓(𝑇), the rate of change in optimal flowering time counting from the start of the season, is caused by 𝑓 declining in the denominator; as 𝑓 gets sufficiently smaller than 1, the subtraction −1/𝑓(𝑇) increases. The 𝑡_2_(𝑇) curvature intensifies even more beyond the range of 𝑇 we observed across our source sites and mathematically has a vertical asymptote (*i.e.* infinite drop in 𝑡_2_). However, this unrealistic extreme limit is biologically irrelevant because the asymptote occurs when 𝑓, maximum relative growth rate, is 0. We expect that the decline of 𝑓(𝑇) will indeed have a biological lower limit, as in the logistic function we fitted. The estimated ranges of 𝑓 (see insets in Fig. 3a and b) roughly corroborate various reports of relative growth rates in the literature (< 0.002∼0.26 · 𝑑𝑎𝑦^–1^ for *Lythrum salicaria* (44–47), and < 0.007∼0.12 · 𝑑𝑎𝑦^–1^ for *Solidago altissima* (48–50), a congeneric species for which reports were available).

### Shifts in spring onset and growing season length together drive accelerating advancement in optimal flowering time

The curvature of 𝑡_2_ = 𝑇 − 1/𝑓(𝑇) represents optimal flowering times when the beginning of the growing season is held equal for all populations (as in our common-garden experiments) by counting 𝑡_2_ values from a common starting line. This allowed us to isolate the role of genetic differences in the time taken to flowering from the beginning of the growing season. However, in nature, the start of the growing season covaries strongly with growing season length (*i.e.,* earlier spring ≈ longer growing season; SI Appendix Fig. S2). Combining variations in growing season length and variations in start of spring at the source population sites tilts the 𝑡_2_ curve clockwise, to give the expected flowering times *in situ* (Fig. 4, solid colored curves).

**Figure 4.**
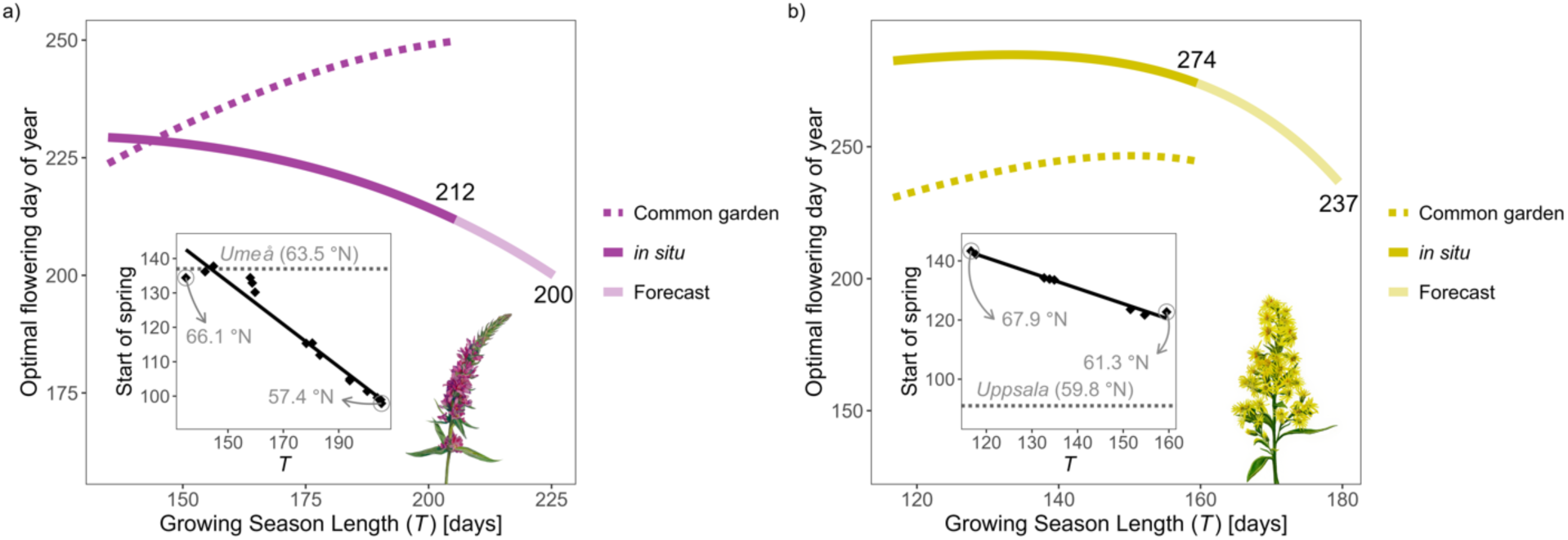
Estimates of *in situ* flowering time across a) *Lythrum* and b) *Solidago* populations, and predictions beyond the observed range of growing season lengths. Dotted gray line in each inset shows the start of spring at the common-garden sites of the respective experiments (Umeå for *Lythrum*, Uppsala for *Solidago*), in the experiment year. Black points are mean *in situ* start of spring against growing season length between 1961 and year of plant collection at the source sites (1997 for *Lythrum*, 2003 for *Solidago*), and black lines are linear regressions. Dotted colored curves in the main plots are fitted optimal flowering (𝑡_2_) predictions across source sites in the common-gardens (flowering time expressed as number of days after the start of the growing season), identical to Fig. 3. Colored solid curves show *in situ* optimal flowering (expressed as day of the year) across *T*, which are summations of the *in situ* start of spring regression lines and the 𝑡_2_ curves. Lighter curves are extrapolations for a further 20-day increase in 𝑇 under the fitted functions, with predicted optimal calendar day of flowering labeled. *Illustrations courtesy of Becca Huggins*.

The tilted curves resulting from considering both the start of spring and growing season length predict the optimal calendar day of flowering *in situ* across the latitudinal gradient. They also predict optimal flowering time in a single locality under climate change, assuming that the start of spring and growing season length change together (as supported by historical data from these sites; SI Appendix Fig. S3). These curves of flowering time advancement have important implications. First, at higher latitudes (*e.g.,* 𝑇 ≲ 150 for *Lythrum*; ≲ 130 for *Solidago*), which represent growing season lengths at less advanced stages of climate change in a space-for-time perspective, the rate of change in optimal flowering time is slow, near stasis. The specific slope of this optimality change—and the duration of relative stasis—in other species will depend on their 𝑓(𝑇) function and the slope of start of spring against growing season length (Fig. 4, insets) of their environments. Second, beyond some critical growing season length 𝑇, the change in optimal flowering time accelerates. The threshold of 𝑇 for this acceleration will again depend on the species, driven by a decreasing 𝑓 in the denominator. However, because 𝑓 is in the denominator, this acceleration should occur in any species as long as 𝑓 continues to decrease with longer 𝑇. Direct measurements of the lower asymptotic limits of 𝑓(𝑇) will clarify the limits of flowering time advancement beyond the acceleration phase.

The nonlinear shape of the tilted 𝑡_2_(𝑇) (Fig. 4) could explain the observed variability in flowering phenological shifts across different systems. Different species, each with unique 𝑓(𝑇) functions, in environments with different correlations between start of spring and growing season length, are expected to exhibit a range of phenological responses from delays to relative stasis to gradual and rapid rates of advancement.

Finally, we extrapolated the tested function by extending 𝑇 by 20 days—which is the average observed increase in 𝑇 across sites since 1961 to present (SI Appendix Fig. S3)—to simulate continued trends under climate change (1, 2, 51). If the current relationship between start of spring and 𝑇 (Fig. 4 insets) stays constant, this extrapolation predicts nonlinear advancements in optimal flowering time for the two species that are rapid for both but quite different in magnitude (12 days for *Lythrum* and 37 days for *Solidago*), just beyond the range of growing season lengths observed at the sites of our study populations (Fig. 4). Assuming that 𝑓(𝑇) continues to follow the fitted logistic functions, extrapolating 𝑇 by 20 days results in 𝑓 = 0.006 for *Lythrum* and 𝑓 = 0.008 for *Solidago*. These values are within the ranges of comparable rates in the literature mentioned above, indicating that these forecast predictions remain biologically realistic. For comparison, if a linear correlation assumption were used to forecast by taking the derivative of the tilted 𝑡_2_ curves (-0.50 for *Lythrum* and -1.0 for *Solidago*) at the maximum observed 𝑇 for each species, it would underestimate flowering time advancement by 2 days for *Lythrum* (10-day advancement) but by 17 days for *Solidago* (20-day advancement), compared to our nonlinear forecast for a 20-day extension of 𝑇. Note that these are conservative error estimates compared to using a linear correlation across the entire observed *T* range.

## DISCUSSION

The growing season is a critical window for all essential annual life-history activities. Despite variations among species in specific activities and their impact on fitness, annual scheduling within limited favorable conditions is a unifying process that can be useful for explaining existing geographical variations in phenology and predicting shifts under climate change. Based on the theory of optimal energy allocation scheduling, we predicted a nonlinear relationship between growing season length and optimal flowering time. This relationship ultimately leads to a sharp fall in optimal flowering time due to an inverse term involving maximum relative growth rate, a key life-history trait. Two independent common-garden experiments validated this functional relationship. What follows is a quantitatively testable prediction that the combined effect of spring advancement and growing season expansion should generate a spectrum of responses from relative stasis to gradual and rapid advancement under the conditions predicted with climate change.

While growing season expansion is a globally reported phenomenon under climate change, the rate and symmetry of that expansion varies geographically (1, 2, 52–55). In particular, the contribution of autumn delay to growing season dynamics is much less investigated than that of spring advancement (56)(but see: (57–59)). In lower latitudes, one further complicating feature is the increase in summer dry periods that may effectively reduce or break up the growing season, even if the boundaries of the growing season defined by temperature thresholds might be expanding (60–62). The additional physiological costs of increased mid-season droughts on optimal flowering time, for instance via effects on maximum relative growth rate 𝑓, needs further investigation. Optimal energy allocation models are well suited for such investigations. Importantly, due to species differences in responses to climatic conditions, the exact climatic boundaries are specific to a given system; the 5-day 5℃ boundary condition we used here is broadly applicable and common, but not a universal condition. While we focused on isolating the general role of growing season length as it is a universal variable that is underexplored, tailoring the model for predictions in specific systems should involve careful consideration of any known climatic boundaries for seasonal growth.

Iwasa & Cohen’s optimization solution predicted an outsized role of maximum relative growth rate 𝑓 in mediating the relationship between growing season length and optimal flowering time. Other canonical life-history traits such as survival, age, size, and reproductive investment were also in the initial model, but cancelled out in the optimization. Intriguingly, this corroborates a long-standing tenet of plant life history theory, that relative growth rate is a comprehensive trait through which many important fitness-related traits can be explained (15, 32, 40, 42). In our study, 𝑓 was a free parameter that we estimated because it is not a standard trait that is easy to measure in plants. However, 𝑓 can be inferred from individual growth curves, *e.g.* by fitting a logistic function to growth timeseries and computing the peak of its derivative (rate of acceleration). There is still the choice of which measurable part of the plant would most accurately reflect the 𝑓 parameter, and this choice will depend on the species. The clinal variation shape of 𝑓 along latitudinal (or growing season length) gradients will also vary between systems. Here we chose a logistic function that is biologically realistic and flexible. The shape parameters that govern the slope and curvature of the function 𝑓(𝑇) = 𝑎⁄1 + 𝑒^−𝑏(𝑇+𝑐)^ have strong influences on the resulting curvature of 𝑡_2_ = 𝑇 − 1/𝑓(𝑇). While 𝑓 might indeed be a conserved trait in some species, we posit this would be an exception rather than the rule, as growth rate is a well-established axis of trade-off with other canonical life-history traits (15, 32, 40, 42). Any variation in 𝑓 with growing season length would cause a nonlinear relationship between growing season length and optimal flowering time in a species, which can help explain and predict flowering time shifts under changing seasonality.

Common-garden experiments are ideal for our validation because they eliminate the impact of unmeasured *in situ* environmental differences. However, several microhabitat features or ecological interactions can alter the trajectory of evolution *in situ* from what we have predicted based on climatic selection on optimal phenology. For instance, by incurring extra cost functions, herbivory (63) and density-dependence (64) might alter the control function 𝜇(𝑡)(0 ≤ 𝑡 ≤ 𝑇), *i.e.,* how the ratio of the growth of the vegetative production part to its maximum rate changes through time (Box 1). In other words, herbivory and density-dependence may dampen the growth of the production part of the plant, and possibly require additional compensations such as maintenance costs, for the part of the season when they are applicable. As all solutions of the optimization directly or indirectly hinge on this control function, such influences could change the optimal switching times including flowering time. Other factors such as interactions with pollinators and seed predators may also influence the control function 𝜇(𝑡) for a subset of 𝑡 (65), and adjust optimal solutions. Beyond altering optimal predictions, temporal variations in such biotic factors and numerous abiotic ones (*e.g.,* soil moisture) can cause plastic variation in phenology. Here, we focused on life history adaptation to seasonal time pressure over many generations, assuming that such other factors are reasonably similar or that they vary unsystematically across compared populations. In both species independently, we found evidence for considerable genetic differentiation in flowering time between populations that follows an optimal prediction.

What determines evolutionarily optimal phenological timing in changing seasonal environments involves two key complexities. First, the life-history processes underlying an individual’s life cycle scheduling are interconnected through covariances and trade-offs. The assumption that earlier springs should drive earlier phenology–a common postulate in the field–oversimplifies the relationship between seasonality and phenology to an imprecise linear one. It suggests that phenology is primarily a product of year-to-year plastic responses, unburdened by life-history costs. This common perspective falls short in explaining the observed variability in phenological shifts (7) or predicting potential limits on future trends. Life history theory emphasizes long-term selection and fitness optimization beyond immediate plastic reactions to environmental changes. To fully confront that phenology is a life history phenomenon, we must consider how interconnected life-history processes reschedule together as the start and duration of the growing season changes. Second, while the onset and duration of the season experienced iteratively over an individual’s lifetime are related, they impose distinct and synergistic influences on optimal timing. Fortunately, optimal energy allocation models offer a powerful framework to address both complexities in a mathematically explicit and customizable manner. Our extended Iwasa-Cohen hypothesis (Eq. 2) predicted a nonlinear pattern of optimal flowering time variation across growing season length gradients—one that would have remained invisible under the assumption that optimal phenology is solely a function of when spring begins. Ultimately, we advocate developing *a priori* quantitative hypotheses grounded in life history theory to move beyond correlative conclusions. This approach can clarify our understanding of how seasonal time pressures select for timing patterns in general and support systematic predictions that can be tested and compared across different systems.

## METHODS

### Optimal flowering time

We base our study on the optimal scheduling model of perennial plants initially developed by Iwasa & Cohen (29) (Box 1). With 28 (life-history and environmental) parameters, the original model can be tailored to explore several representative classes of plant life histories. We focus on a major class that also mirrors our empirical test systems, namely plants that: (1) grow at the beginning of each season using reserves from the preceding year as well as current-year photosynthesis, and (2) exhibit gradual (non-instantaneous) re-growth of photosynthetic parts at the onset of each season. Briefly, the model breaks down each year into three phases, highlighting key moments when a plant should switch from using stored resources to fully relying on current photosynthetic growth (𝑡_1_), and then to investing in reproduction and storage (𝑡_2_), with the latter marking the timing of first flower. Given the length of the growing season experienced each year, a combination of yearly and between-year optimizations determines the solutions (optimal values) of the two critical switching times.

#### Box 1.

**Summary of the optimal energy allocation model for perennial flowering time**

##### Model overview

An individual perennial plant consists of a “production” part (𝐹) and a “storage” part (𝑆). The former comprises vegetative organs that drive energy generation and growth, *i.e.* leaves, stems, and roots, but not trunks or larger branches. The latter encompasses resources stored for use in the following season, as well as allocation to reproductive organs (*i.e.* flowers and fruits). The objective of the model is to optimize the scheduling of energy allocation to maximize lifetime reproductive investment. Scheduling decisions must balance maximizing each year’s reproduction with preserving an optimal amount of resources for future reproduction – thus, the problem becomes one of dynamic optimization.

##### Summary of the dynamics and optimization

Daily net production rate depends on the size of 𝐹, and drives the growth of 𝐹 itself as well as 𝑆. This production rate follows the function 𝑔(𝐹) = 𝑓𝐹/(1 + ℎ𝐹), where 𝑓 is the plant’s inherent maximum relative growth rate, which is realized when the plant is small, and ℎ is a self-limiting coefficient.

While the original theoretical framework explores a spectrum of life history scenarios, from here we specifically trace derivations that reflect both our empirical systems and a typical perennial life history. Namely, there are three distinct phases across time 𝑡 in year 𝑛, bounded by growing season length 𝑇 (Fig. 1): (*1*) 𝐹_𝑛_grows by using both storage 𝑆_𝑛_(𝑡) and current production 𝑔[𝐹_𝑛_(𝑡)]. Given conservation of material, 𝑑𝐹_𝑛_⁄𝑑𝑡 = 𝑔[𝐹_𝑛_(𝑡)] − 𝑑𝑆_𝑛_⁄𝑑𝑡. (*2*) Storage is depleted, marking the first critical switching point 𝑡_1_. From this point, 𝐹_𝑛_ continues to grow using production at a rate bounded by 0 ≤ 𝑑𝐹_𝑛_⁄𝑑𝑡 ≤ 𝑎𝐹_𝑛_ + 𝑏, where 𝑎 and 𝑏 are limiting constants. (3) The second critical switching point 𝑡_2_ occurs when vegetative growth stops, and storage growth 𝑑𝑆_𝑛_⁄𝑑𝑡 begins.

Both switching points follow “bang-bang control”, meaning an immediate as opposed to a gradual switch, which is a canonical result of evolutionary optimal theory supported by many empirical studies (19, 30, 66, 67). While the model does not distinguish between the onsets of flowering, fruiting, or storage *sensu stricto* within the third phase, 𝑡_2_ marks the end of vegetative growth and the beginning of reproductive investment. Thus we treat 𝑡_2_ as flowering time (*cf.* (27, 68)).

At the end of the growing season 𝑡 = 𝑇, any remaining 𝐹_𝑛_ is discarded (*e.g.* leaf shedding), to be reconstructed from storage in the next season. In the beginning of growing season 𝑛 + 1, the size of the storage is 𝑆_𝑛+1_(0) = 𝛾[𝑆_𝑛_(𝑇) − 𝑅_𝑛_], where 𝛾 is storage efficiency and 𝑅_𝑛_is reproductive investment in year 𝑛.

The overall objective is to maximize lifetime reproductive investment 𝜙 over 𝑛 years, *i.e.* find the strategy in which 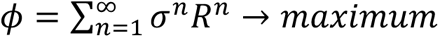, where 𝜎 is annual survival. This dynamic problem is necessarily divided into ‘within-season’ and ‘between-season’ subproblems, where the solution to the former informs the latter as in a mathematical recursion. The first is solved using Pontryagin’s Maximum Principle from optimal control theory. Here the variable 𝜇(𝑡)(0 ≤ 𝑡 ≤ 𝑇) is the focal control variable to be optimized, which is the ratio of the growth rate of the production part to its maximum, such that 𝑑𝐹_𝑛_⁄𝑑𝑡 = 𝜇(𝑡)(𝑎𝐹 + 𝑏). The solution involves maximizing the Hamiltonian 𝐻(𝐹, 𝑆, 𝜆_𝐹_, 𝜆_𝑆_, 𝜇) = 𝜆_𝐹_𝜇(𝑎𝐹 + 𝑏) + 𝜆_𝑆_[𝑔(𝐹) − 𝜇(𝑎𝐹 + 𝑏)] , where 𝜆_𝐹_, 𝜆_𝑆_ are co-state variables of 𝐹 and 𝑆. In a simple sense, to find the optimal 𝑡_2_ is to find when 𝜇(𝑡) should first equal 0. The second problem of optimally dividing each year’s 𝑆_𝑛_(𝑇) into 𝑅_𝑛_ and storage, which dictates the optimal schedule for year 𝑛 + 1 and so on, is solved using another mathematical technique called dynamic programming.

The solution of the combined subproblems that we explore in this study is the optimal flowering time as a function of growing season length, which is 𝑡_2_ = 𝑇 − 1/𝑓. This solution is notably independent of the plant’s age (𝑛) or size. We remark that 𝑓 itself has been found to be related to 𝑇 in many systems. This modifies the solution to 𝑡_2_ = 𝑇 − 1/𝑓(𝑇), giving an inverse function as opposed to linear, whose downward curvature accelerates as 𝑇 increases.

Note that the optimality solutions represent long-term evolutionary direction, as opposed to momentary, or plastic, responses to short-term variation in season length. We thus assume that in our empirical systems, each population *in situ* is at or near its evolutionary equilibrium at the time of sampling, and that differences among the populations in the experiments reflect differences in these equilibria. Additionally, we assume that by sampling across many source populations (15 for *Lythrum* and 8 for *Solidago*), any differences in the effects of ecological interactions *in situ,* such as density-dependence or herbivory, are minimized as random noise, allowing us to detect signals of the more basal, internal life history optimization pressures imposed by climate seasonality. Nonetheless, the comparison of flowering time among populations was made using common-garden experiments, and thus differences among populations are assumed to reflect genetic differences.

### Growing season lengths, and start of spring in source populations

The climatological, or meteorological, growing season is defined as the duration within the year in which plant growth can theoretically take place (1, 53). Species vary in the relevant thermal conditions that bound their growing seasons. However, a conventional metric that is widely deemed physiologically favorable (for growth) and safe (from frost) for perennial plants, especially in mid-to high-latitudes in the Northern Hemisphere, is when the threshold of 5°C daily mean air temperature (>5°C for start of spring and <5°C for start of autumn) for a sustained period (≥ 5 consecutive days) is crossed (1, 52, 53, 69–71). We obtained historical daily mean air temperature data for each source population’s site from the high-resolution gridded PTHBV database generated by the Swedish Meteorological and Hydrological Institute (www.smhi.se). Date ranges of the obtained data spanned from 1961 (earliest year in the database) to years when plants were collected for the respective common-garden experiments (1997 for *Lythrum*, and 2003 for *Solidago* populations). We calculated the start and end of the growing season for each population in each year, with the difference representing the length of the growing season, and subsequently calculated the average growing season length for each population across its year range.

### Lythrum salicaria *common-garden experiment*

Purple loosestrife, *Lythrum salicaria* (72), is a perennial herb native to Eurasia that has been introduced widely to North America, and Australia. In Europe, it is found across the Mediterranean to northern Fennoscandia (36-67°N). It occurs in a variety of wetland, lake- and seashore, riparian, and fen habitats. We used data collected and published in (43), which contains more detailed information about the sampling protocol and experimental design. Fifteen populations across a wide latitudinal gradient of Sweden (57-66°N) were chosen, and seeds were collected from a minimum of 200 individuals per population (SI Appendix fig. S1b). In May 1997, seeds from 28-62 randomly chosen maternal plants from each source population were planted in the greenhouse at Umeå University in northern Sweden (63°49’N). The plants were grown under a photoperiod of 16h, and 18°C-day and 8°C-night temperature cycles. In June 1997, the plants were repotted and transferred to the outdoor experimental garden in a fully randomized design (each maternal family represented by a single plant). In 1999, the date that the first flower opened was recorded for each plant. For our analyses, we converted flowering date to a numerical date relative to the start of the growing season.

### Solidago virgaurea *common-garden experiment*

The European goldenrod or woundwort, *Solidago virgaurea*, is a perennial herb widely distributed across Europe, northern Africa, and Asia (72). In Sweden, it occurs in forest and meadow vegetation and reaches into the alpine zone of the Scandinavian mountain range. In the first year, the plant forms a leaf rosette. In the following years, the plant can flower or be vegetative, with flowering individuals producing one or several vertical inflorescences. To assess among-population variation in flowering time, and the association between flowering start and length of the growing season at the site of origin, we conducted a common-garden experiment in the Botanical Garden of Uppsala University (59°50’N). For the present analysis, we used data on flowering time of eight boreal populations located along a latitudinal gradient in Sweden (61-68°N; SI Appendix Fig. S1b). Seeds for the experiment were collected between late August and late September 2003. About two months after collection, we planted seeds in 11×11×12 cm plastic pots in an 80:20 mixture of commercial potting soil (“Weibulls yrkesplantjord”, Weibulls, Hammenhög, Sweden) and LECA (Light Expanded Clay Aggregate, 2-6 mm in diameter, AB Svensk Leca, Linköping, Sweden). Pots were arranged in a randomized block design with 25 blocks. In each block, each population was represented by one plant from a different maternal family. We planted five seeds per pot and emerging plants were thinned to one plant per pot the following spring. In 2005, the great majority of plants in the experiment had reached the flowering stage (median [range], 96% [88-100%]; N = 8 populations). We monitored flowering twice a week once the first flower buds were observed, and for each plant the first day of flowering was recorded. Prior to analysis, we converted flowering date to a numerical date relative to the start of the growing season.

### Statistical analyses

We tested statistical support for the Iwasa-Cohen model using non-linear Bayesian regression models, and compared non-linear models to models assuming constant relative growth rate, 𝑓, using cross validation. Our non-linear models comprised a functional relationship between relative growth rate 𝑓 across growing season length 𝑇 ( 𝑓(𝑇) ), as well as the optimized relationship between flowering time (𝑡_2_) and 𝑇 (Eq. 2). We specified a flexible logistic function of relative growth rate 𝑓 across growing season length 𝑇 with three shape parameters, of the form 𝑓(𝑇) = 𝑎⁄1 + 𝑒^−𝑏(𝑇+𝑐)^ (Fig. 2, inset). Note that this function includes the possibility of no change in 𝑓 across 𝑇, namely when 𝑏 = 0. Then, we fitted the function 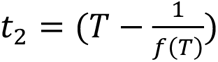 to the data. In constant 𝑓 models, we explored the linear (slope constrained to 1), intercept-only relationship between 𝑡_2_and 𝑇. We fitted all models in a Bayesian regression framework using the *brms* package of R (R version 4.3.2; (73)). In all models, we assumed a Gaussian distribution of 𝑡_2_ (SI Appendix Text S1). We ran models across four chains for a total of 4000 iterations, with 2000 warm up iterations, and assessed model convergence using 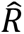 values (measure of chain mixing). For constant 𝑓 models, we used weakly informative normal priors. In non-linear models, we used highly regularized, bounded normal priors, to avoid fitting errors due to the asymptotes implied by the inverse term 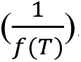. We explored prior bounds for the non-linear models using visual prior predictive analysis. For full model structures and prior specifications, please refer to SI Appendix Text S1. Following model fitting, we compared the predictive performance of linear and non-linear models using leave-one-out cross validation implemented in the *loo* package (74). We used the differences in expected log pointwise predictive density ( 𝑒𝑙𝑝𝑑) to compare model predictive performance. Finally, we assessed model parameter uncertainty at the 99% level, and predictions also included population-level variance 𝜎 calculated at the 50% quantile level.

## ACKNOWLEGEMENTS

We thank Stephen C. Stearns for feedback and comments on a draft. We thank Becca Huggins for the illustrations of *Lythrum salicaria* and *Solidago virgaurea* inflorescences in Figures 3 and 4. This work was financially supported by the European Union’s Horizon 2020 research and innovation programme under the Marie Skłodowska-Curie grant agreement No 101030973 (“Cycles of Life”) to JSP, Horizon Europe Marie Skłodowska-Curie grant agreement No 101067850 (“ClimRes”) to JJ, and a grant from the Swedish Research Council (2023–05270) to JÅ.

