## Supplementary material for "Testing a general theory for flowering time shift as a function of growing season length": SI Appendix

**This PDF file includes:**

Figures S1 to S3

Supporting Text S1

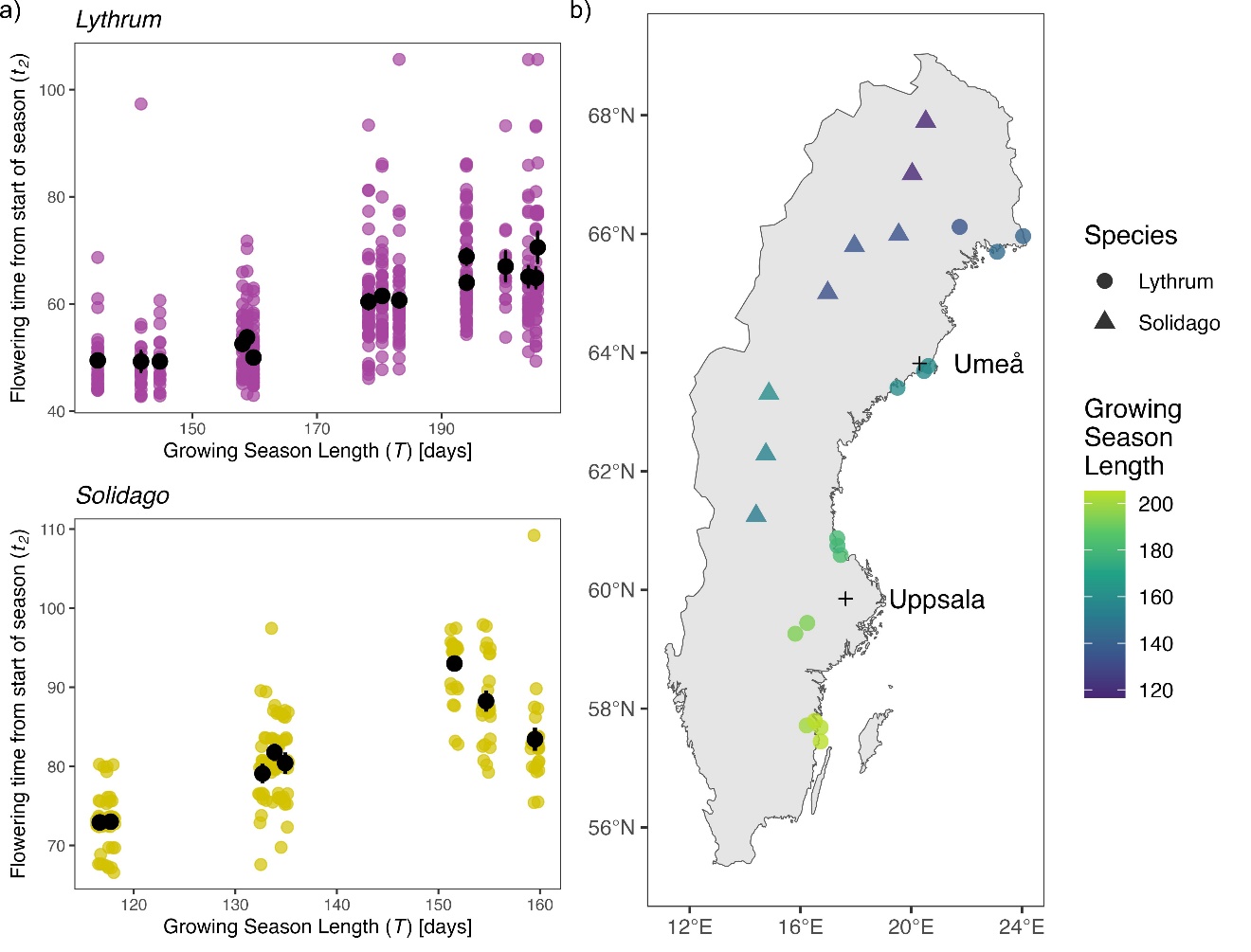

**Figure S1. Flowering time data and source population sites.** a) Colored points show individual plant flowering times (flowering initiation) measured from the start of spring ($t_{2}$) in the respective common garden experiment location in the experiment year (Umeå in 1999 for *Lythrum*, and Uppsala in 2005 for *Solidago),* plotted against growing season length ($T$) of source population site. $T$ is the average growing season length from 1961 to collection year (1997 for *Lythrum* and 2003 for *Solidago*), measured as duration between the first and last time the threshold of 5 consecutive days of 5ºC daily mean temperature is crossed. b) Circle and triangle points show locations of *Lythrum* and *Solidago* source populations, respectively, from which seeds for the common garden experiments were collected. Colors are growing season lengths ($T$) at each locality.

**Supporting Information Text**

Supporting Information Text S1. Full model structure and prior distribution specifications for the Bayesian regression models.

**Constant** $\boldsymbol{f}$ **(original Iwasa-Cohen) model structure (both *Lythrum* and *Solidago*)**

$$t_{2} \sim Normal\left( \mu, \sigma\right)$$

$$\mu_{i}=T_{i}-1/f$$

Priors

$$f \sim Normal\left( 0, 0.1 \right)$$

$\sigma\sim Normal(0, 5)$, for $0<\sigma$

**Extended Iwasa-Cohen (variable** $\boldsymbol{f}$**) model structure: *Lythrum***

$$t_{2} \sim Normal\left( \mu, \sigma\right)$$

$$\mu_{i}=T_{i}-\frac{1}{a/{1+e^{-b(T+c)}}}$$

Priors

$a \sim Normal\left( 0.05, 1 \right)$, for $0.0001<a<1$

$b \sim Normal\left( -0.05, 1 \right)$, for $-1<b<-0.0001$

$c \sim Normal\left( -150, 10 \right)$, for $-200<c<-100$

$\sigma\sim Normal(0, 5)$, for $0<\sigma$

**Extended Iwasa-Cohen (variable** $\boldsymbol{f}$**) model structure: *Solidago***

$$t_{2} \sim Normal\left( \mu,\sigma\right)$$

$$\mu_{i}=T_{i}-\frac{1}{a/{1+e^{-b(T+c)}}}$$

Priors

$a \sim Normal\left( 0.5, 0.75 \right)$, for $0.0001<a<1$

$b \sim Normal\left( -0.5, 0.75 \right)$, for $-1<b<-0.0001$

$c \sim Normal\left( -150, 10 \right)$, for $-200<c<-100$

$\sigma\sim Normal(0, 5)$, for $0<\sigma$

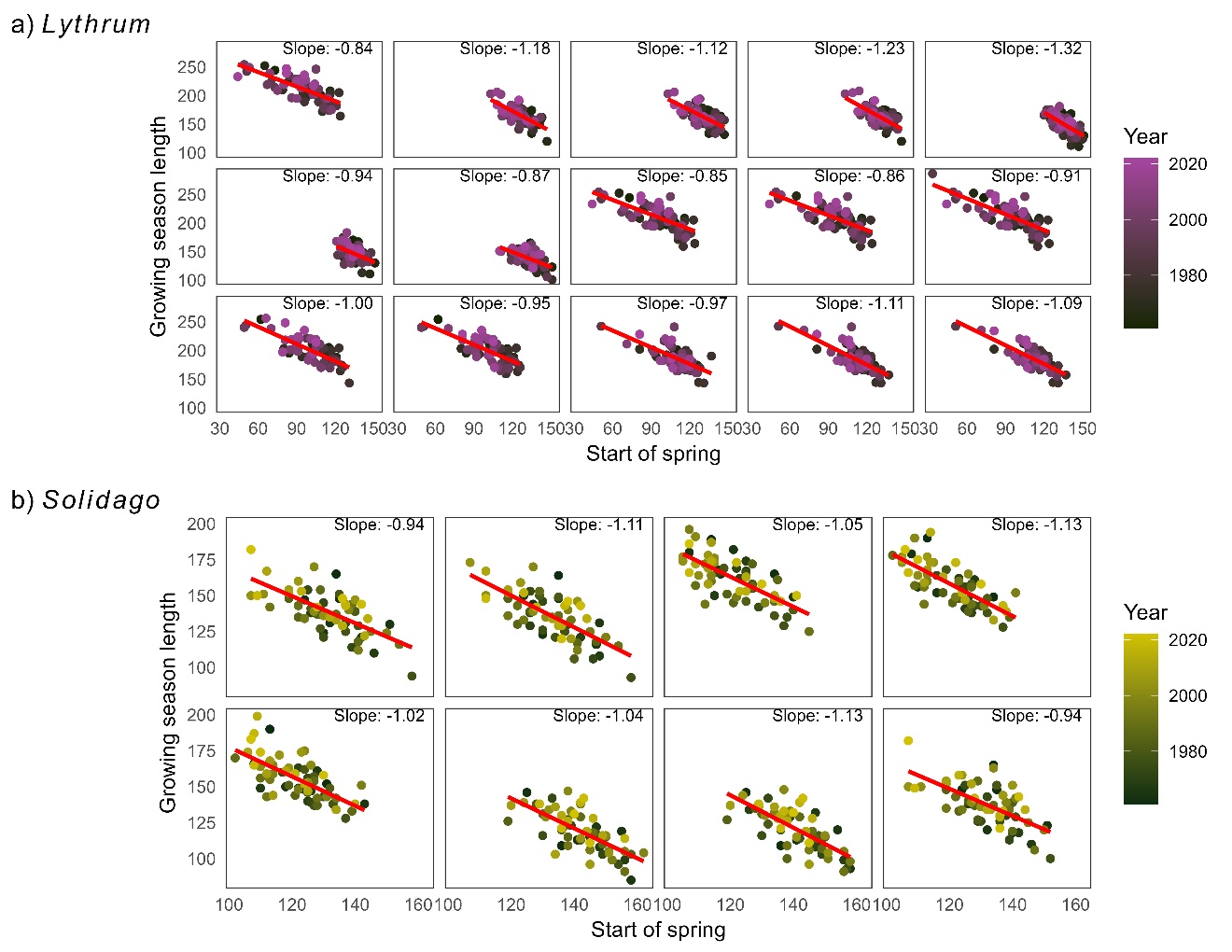
**Figure S2. Association between growing season length and start of spring at a) *Lythrum* and b) *Solidago* source sites from 1961 to 2022.** Each panel represents a source site, with colored points corresponding to year. Lines and slope values show linear regression in each site.

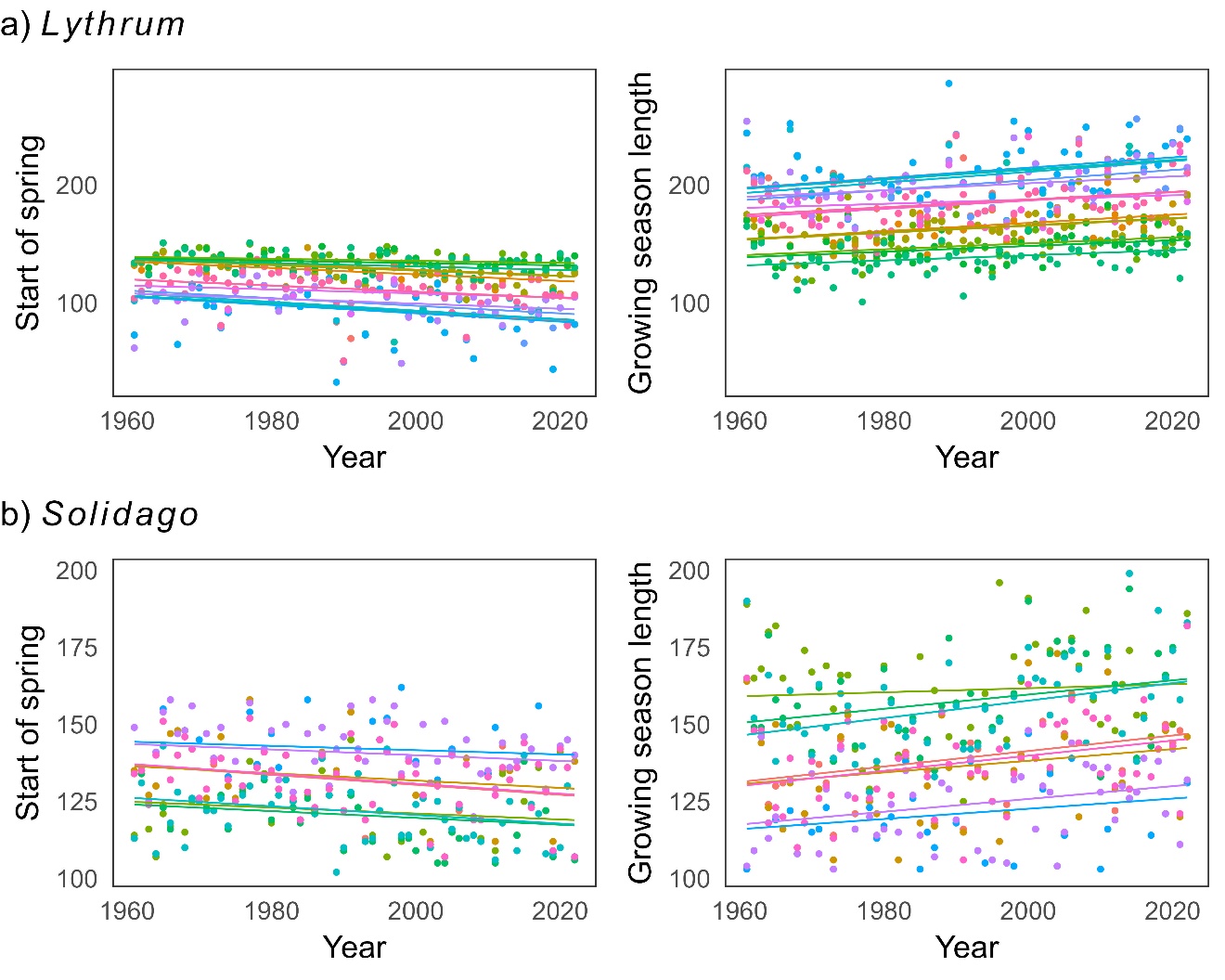

**Figure S3. Historical trends in start of spring and growing season length in a) *Lythrum* and b) *Solidago* source sites.** Colors correspond to source sites (15 for *Lythrum* and 8 for *Solidago*), with points being metrics per year and lines being linear regressions through them.
